# Underground Guardians: How Collagen and Chitin Amendments Shape Soil Microbiome Structure and Function for *Meloidogyne enterolobii* Control

**DOI:** 10.1101/2024.06.18.599572

**Authors:** Josephine Pasche, Roshni Sawlani, Victor Hugo Buttrós, Johan Desaeger, Karen Garret, Samuel J. Martins

## Abstract

The emergence of the Guava Root-Knot Nematode (*Meloidogyne enterolobii*) poses a significant threat to tomato yields globally. This study aimed to evaluate the impact of collagen and chitin soil amendments on soil microbial composition and function (fungal and bacterial communities), and their effects on tomato plant health and *M. enterolobii* infection under standard (5,000 eggs plant^-1^) and high (50,000 eggs plant^-1^) inoculum pressure. Conducted in a greenhouse setting, the study investigated the effectiveness of these amendments in nurturing beneficial microbial communities across both native and agricultural soils. Both collagen and chitin were effective in reducing nematode egg counts up to 66% and 84% under standard and high inoculum pressure, respectively and enhance plant health parameters (biomass and chlorophyll content). Moreover, a microbiome shift led to an increase in bacterial (*Kitasatospora, Bacillus, and Streptomyces*) and fungal (*Phialemonium*) genera, known for their chitinase, collagenase, and plant-parasitic nematode control. Among the microbes, *Streptomyces* spp. were found among the core microbiome and associated with a lower disease incidence assessed through a phenotype-OTU network analysis (PhONA). Under standard inoculum a higher metabolite expression was observed with the amino acid class being the majority among the metabolite groups. The findings highlight the potential of collagen and chitin to mitigate *Meloidogyne enterolobii* infection by fostering beneficial soil microbial communities.

## Introduction

The root-knot nematode, *Meloidogyne enterolobii,* is a global threat to a broad range of plants, including a wide variety of tomato cultivars, with the damage from this species alone causing up to 65% yield loss (Cetintas et al., 2007). The use of chemical controls such as nematicides may not be an economically viable management tool, due to high costs and negative impacts on the environment and human health (Tuncsoy, 2021). These issues lead to the increasing need for alternatives to chemical control. The use of more sustainable management approaches, such as the use of beneficial microbes have been shown to be effective in promoting plant health and suppressing plant diseases (Martins et al., 2023). Beneficial microbes have also been shown to be effective specifically in controlling infection by *Meloidogyne* spp. (Pasche et al., 2023; Rashidifard et al., 2022; Vieira de Carvalho Júnior et al., 2022).

Microbes can lead to an antagonistic effect against nematodes, and soil amendments may be used to favor beneficial microbes over others (Pasche et al., 2023). Approximately 80% of the nematode cuticle is composed of collagen, and with addition of collagen to the soil, bacteria that produce collagenases are likely to accumulate (Page et al., 2014; Raina et al., 2019). The collagenase activity may damage the nematode cuticle, resulting in decreased nematode populations in the roots of tomato plants (Galper et al., 1990; Galper et al., 1991). Likewise, chitin is an essential component in the eggs of nematodes, and chitinase producing microorganisms may have an impact on the presence of nematode infections. Tian et al. (2000) isolated chitinolytic bacteria by adding chitin (1% w/w) to the soil of soybean plants in a greenhouse trial, allowing the bacteria to inhabit the soil, and then isolating with selective media. These isolates were tested *in vivo* and effectively controlled nematode (*H. glycines*) infections. In this study, we hypothesized that soil amendment with collagen and chitin will alter soil microbial assemblages to favor beneficial microbes, decrease nematode infection, and contribute to overall plant health.

The objectives of this study are as follows.1) Determine the effects of collagen and chitin amendment on the soil microbial composition, analyzing the fungal and bacterial microbiome networks and whether microbes that produce collagenase and chitinase are favored. 2) Conduct a second experiment utilizing heightened inoculum pressure to assess the core microbiome constituents that persist under such stress conditions. 3) Assess the enzymatic activity in the soil microbiome in response to the addition of the soil amendments. 4) Assess the possible change in soil health, tomato plant health, and severity of guava root-knot nematode infection with the addition of collagen and chitin to the soil.

## Materials and Methods

### Soil collection

Two soils were used in this experiment, a native soil and an agricultural soil. The agricultural soil is from an organic tomato field, collected at the University of Florida Gulf Coast Research and Education Center, Wimauma, FL field (39.6 altitude, 27°45’23’’ N and 82°13’29’’ W). We collected the native soil from an uncultivated area covered in native foliage approximately 30 ft from the tomato field at the University of Florida Gulf Coast Research and Education Center, Wimauma, FL (39.6m altitude, 27°45’23.7168” N and 82°13’27.876” W). The two soils were chosen with proximity so that they are more likely to have similar microbial and physical composition.

Soils were mixed with 50% potting soil to assist in moisture retention, given the sandy soil texture of Florida soils. We mixed the in a cement mixer to ensure homogeneity between the amendments and the soil before filling the pots. Fresh soil was collected before starting each experiment, with a three-month interval between collections. Physical-chemical analysis of the soils was then conducted by the University of Florida Extension Soils Lab (Table 1).

**Table 1.**
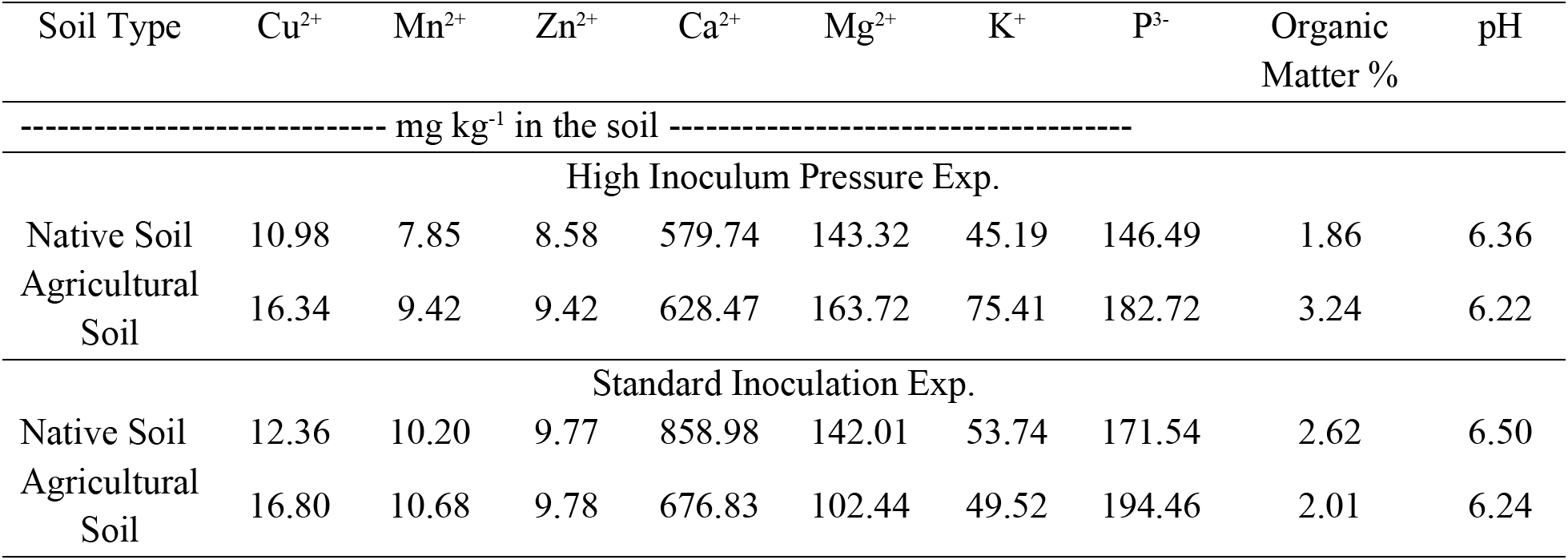
Soil chemical parameters assessment (organic matter content, macro- and micronutrients) for the two types of soils used in this study.

For both experiments, an overall increase in nutrient levels was observed for agricultural soil, with few exceptions.

### Experimental design

Two *in vivo* greenhouse experiments were conducted, focusing on the influence of soil amendments and the increased *M. enterolobii* on plant health and soil microbiome dynamics.

The first experiment was conducted to evaluate the impact of various soil amendments on plant health, focusing particularly on their potential to influence the soil microbiome composition and mitigate disease susceptibility. We determined a standard inoculum of 5,000 eggs per pot based on use in prior literature (Bui & Desaeger, 2021; Brito et al., 2020). This foundational experiment established a baseline understanding of how alterations in the soil environment could affect plant-microbe interactions and, by extension, plant health. Building upon the insights from this initial study, a more rigorous assessment was undertaken to determine the resilience and adaptability of the plant-associated microbiome under heightened inoculum pressure. To achieve this, we conducted a second experiment in which the disease challenge was escalated by introducing *M. enterolobii* eggs at a concentration tenfold greater than that used in the preliminary experiment (50,000 eggs). In past work, it has been shown that an increased inoculum concentration of *Meloidogyne* spp. eggs can be linked to lower plant yield (Ekanayake et al., 1984; Maleita et al., 2012). This strategic amplification aimed to simulate a high pathogen load scenario, enabling the analysis of the core microbiome constituents that persist under such stress conditions and clarifying the microbiome’s role in plant disease resistance mechanisms. This escalated inoculum approach also stress-tests the previously observed disease mitigating effects of soil amendments and uncovers the microbiome’s inherent capacity to buffer plants against intensified disease threats, thereby providing valuable insights into potential microbial-based strategies for enhancing plant resilience.

For each soil type, native soil (N) and agricultural (A), there were 4 treatment groups, including a control, and 8 replicates per treatment following a randomized complete block design. The 4 treatments were the control (C), nematode eggs (N), collagen + nematode eggs (Co), and chitin + nematode eggs (Ch). The high inoculum pressure experiment was conducted with the same treatment groups as previously described with 5 replicates per treatment.

### Treatment application

Five weeks after germination, tomato seedlings (cv. Little Napoli) were planted in 2 L plastic pots in a greenhouse (362lm/ft^2^, 28°C, and 65% rh). We fertilized plants weekly (NPK 20-20-20), and hand watered every 24 hours.

Right before planting, collagen and chitin were mixed in the soil using a cement mixer (as described previously). The same day, directly after transplanting the tomato seedlings, 3 small holes were made in the soil close to the root zone, then a total of 5,000 PPN eggs were applied equally divided among the three holes using a 1000 μL pipette. In the high inoculum pressure experiment, 50,000 PPN eggs were applied to the soil using the same method.

### Assessed plant- and nematode-related variables

At 45 days after planting (60 days under high inoculum pressure), we removed the roots from the pots and separated them from their respective shoots, a rhizosphere sample was taken, and the remaining soil was then gently rinsed off. The plant growth parameters fruit fresh weight (FFW), root fresh weight (RFW), shoot fresh weight (SFW), fruit dry weight (FDW), and shoot dry weight (SDW) were taken. To measure the concentration of chlorophyll, the absorbances of chlorophyll A, chlorophyll B, and carotenoids were measured. A 5 mm leaf disk was cut from the third branch of each sample and placed in a 1.5 mL tube with 1 mL of 80% acetone. The samples were incubated at room temperature for 24 hours, then the absorbances were recorded. The resulting data were analyzed using the formulas described by Lichtenthaler and Wellburn (1983) (Table 2).

**Table 2:** Formulas for the calculation of pigments (Lichtenthaler and Wellburn, 1983) Chlorophyll a 12.7(A_663_) – 2.69(A_646_)

|  |  |
| --- | --- |
| Chlorophyll a | $12.7(A_{663}) - 2.69(A_{646})$ |
| Chlorophyll b | $22.9(A_{646}) - 4.68(A_{663})$ |
| Carotenoids | $(1000 A_{470} - 3.27[\text{Chl a}] - 104 [\text{Chl b}])/227$ |
| Total Chlorophyll | $20.2(A_{646}) + 8.02(A_{663})$ |
| Total pigments | Chlorophyll a + Chlorophyll b + Carotenoids |

Then the gall index for roots was assessed using two indices rated on a 0 to 5 scale, based on the number of egg masses and galls. In the Brito et al. index, 0= 0 egg masses and galls per root system; 1= 1-2 egg masses and galls per root system; 2= 3-10 egg masses and galls per root system, 3= 11-30 egg masses and galls per root system; 4= 31-99 egg masses and galls per root system; and 5= ≥100 egg masses and galls per root system (Brito et al., 2020). In the Hussey & Janssen index, 0= 0% egg masses and galls per root system; 1= Few or trace amounts of galls present in a root system; 2= <25% of the root system is galled, 3= 26-50% of the root system is galled; 4=51-75% of the root system is galled; and 5= >75% of the root system is galled (Hussey & Janssen, 2002).

Nematode eggs were extracted from infected roots using the Hussey and Barker (1973) method modified by Brito et al. (2020). We extracted eggs from the roots using 1% NaOCl. The NaOCl solution was added until the liquid just fully covered the roots. The root-NaOCl mixture was blended on the highest setting for approximately 30 secs, and the blended remains added to a series of sieves (No.200 and No.500 mesh, respectively). The nematode suspension was rinsed with a continual stream of tap water until there was no visible trace of bleach on the sieves. The eggs caught by the No.500 sieve were then resuspended in water to a final concentration of 1,000 eggs mL^-1^.

### Rhizosphere isolation and DNA extraction

Soil samples for 16S and ITS sequencing were extracted from the soil rhizosphere (collected prior to root washing), following the protocol from Simmons et al. (2018). First, 5 g of roots were removed with the soil still attached and placed into 25 mL of epiphyte removal buffer (Simmons et al., 2018). The tubes were then placed in an ultrasonic bath for 10 minutes at room temperature to detach microbes from the roots. Roots were then removed, and the remaining solution was centrifuged for 10 mins on the maximum speed. The buffer was then discarded, and the remaining rhizosphere was used for DNA analysis.

The DNA was extracted from 250 mg of soil using the Zymo Quick-DNA™ Fecal/Soil Microbe Miniprep Kit (Zymo Research, Irvine, CA). DNA purity was assessed using spectrophotometry (NanoDrop model, ThermoFisher, Waltham, MA). DNA was sequenced for ITS and 16S analysis at SeqCenter in Pittsburgh, PA.

### Data analysis and statistical design

A t-test was applied for comparison between pigments, egg number, gall counts, and comparison among biomass parameters, dry shoot weight and root fresh weight. For all analyses, the assumption of normality was checked by Shapiro-Wilk and Kolmogorov-Smirnov tests prior to analysis. The Kruskal-Wallis one-way analysis of variance on ranks (*P<*0.05) was applied for significant means when normality was not met. SigmaPlot® version 14.5 was used for statistical analyses.

SHAMAN was used to process and assign taxa to raw fastq.gz files (Volant et al., 2020). Sequences were clustered based on the operational taxonomic unit (OTU) at a threshold of 97% similarity. Taxonomy assignment for the OTUs was conducted utilizing the SILVA database (McDonald et al., 2012). The relative abundances, Alpha and Inverse Simpson diversity indexes were calculated according to the default parameters of Volant et al. (2020). Raw read data was submitted to the NCBI SRA under the accession number PRJNA1120455. After taxonomic assignment, microbial networks were constructed in R v4.3.1 (R Core Team 2024) using methods adapted from Poudel et al. (2023). Bacterial and fungal OTUs were assessed to determine their co-occurrence with desired plant phenotypes, leveraging a Phenotype-OTU (PhONA) network analysis. To initiate this analysis, associations between bacterial and fungal OTUs and disease severity (the phenotype) were evaluated following infection by *M. enterolobii* in tomato plants cultivated in each of the soil treatments. This association was established by identifying treatments that fell within the bottom quartile of eggs per gram roots. Logistic regression was used to establish the log odds of associations between the top 5% most abundant OTUs and the phenotype, thus facilitating an understanding of how the presence or absence of each OTU correlates with the phenotype. OTUs for which p<0.05 were then represented in a network format using the igraph package in R (Csardi & Nepusz, 2006).

In the OTU-OTU network analysis, positive and negative relationships were identified within the microbial community. A table of OTU relative abundance across the soil samples was analyzed. Spearman’s rank correlations were evaluated pairwise for OTU relative abundances, using a significance threshold of p<0.001. These correlations were used to assemble an adjacency matrix, with the corresponding network representation plotted using igraph, with nodes representing OTUs and edges denoting significant correlations.

The core microbiome was assessed for bacterial and fungal OTUs, with core microbiomes determined comparing the experimental treatments soil type (native vs. agricultural soil) and experiment (high vs. standard inoculation). This assessment was made using the microbiome R package (Lahti & Shetty, 2017). To determine the core microbiome, an Abundance-Occurrence approach was taken, defining the “core” as OTU’s found between groups at a 0.001 abundance detection limit and a 50% prevalence (Neu et al., 2021).

## Results

A slight decrease in gall count was observed when amendments were added to the soil (Table 4).

Regarding egg count, nematode infection was observed to be significantly lower in the native soil with the addition of either soil amendment (Fig. 1A).

**Figure 1.**
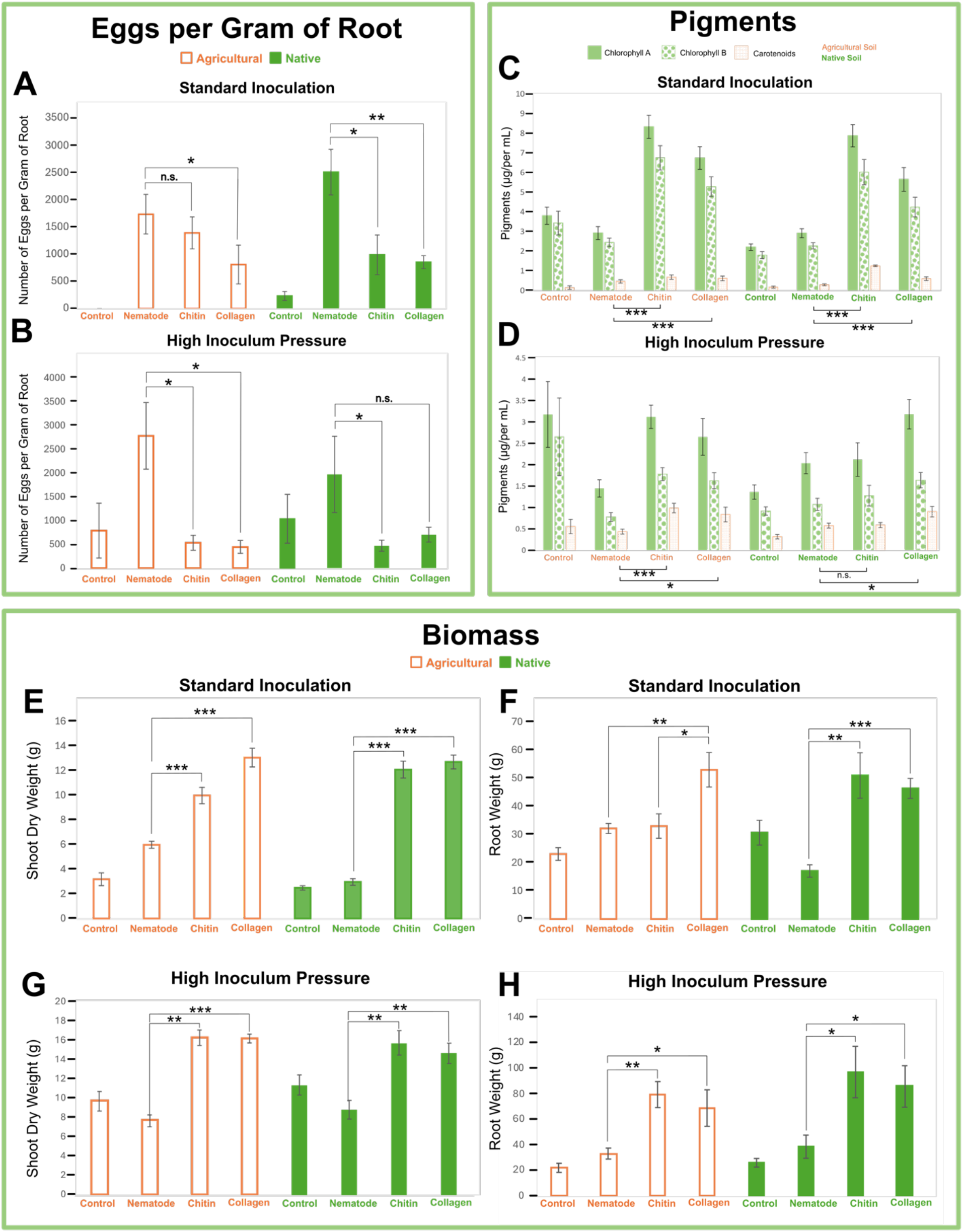
Comparison of plant health parameters and *M. enterolobii* infection in tomato cv. Little Napoli. Number of *M. enterolobii* eggs per gram of root, measured at 45 days (A) and 60 days (B) after inoculation, respectively, showing responses under standard (5,000 eggs) and high inoculation pressure (50,000 eggs). Photosynthetic pigments in leaves under standard (C) and high inoculation (D). Biomass parameters with dry shoot (E) and root (F) weights under standard inoculation; Dry shoot (G) and root (H) weights at high inoculum pressure. *Significant at the 0.05 probability level by Mann-Whitney Rank Sum Test (AB) and Student’s T-test (C-H). Statistical significance is indicated by *p<0.05, **p<0.01, ***p<0.001, n.s. = not significant. Error bars denote ± SE. Sample sizes are 8 replicates for standard inoculation and 5 for high inoculum pressure.

The addition of chitin to the soil reduced egg number by 60% (P<0.01) and collagen reduced eggs per gram of root by 66% (P<0.001). In the agricultural soil, the number of eggs per gram roots decreased in soil amended with collagen (P<0.05); however, we observed a weaker reduction for chitin amended soil (P=0.67).

Under high inoculum pressure, the addition of chitin and collagen in the agricultural soil led to a significant reduction in the number of eggs per gram of root by 81% and 84%, respectively (P<0.01) (Fig. 1B). In native soil, the amendments also resulted in a decrease in the number of eggs, with chitin-amended soils showing a 76% (P=0.03) reduction and collagen-amended soils showing a 64% reduction (P=0.15).

Chlorophyll content also increased, indicating strong evidence of improved plant vigor, with collagen and chitin treatments overall boosting chlorophyll by 1-fold and 1.7-fold, respectively (P<0.001) (Fig. 1C). Similar trends were observed under high inoculum pressure, with amended agricultural soil showing increases in pigment concentration (Fig. 1D). However, in native soil, the chitin group did not significantly increase pigment content compared to the infected control. Furthermore, tomato plants displayed a lower overall pigment concentration compared to the standard inoculation experiment.

With the addition of chitin in the agricultural soil, dry shoot weight increased by 66.72% (P<0.01) when compared to the nematode-infected control (Fig. 1E). Similarly, adding collagen to the agricultural soil yielded a 1.2-fold increase in dry weight (P<0.001). In native soil, the addition of chitin and collagen yielded a 3.1-fold (P<0.01) and a 3.3-fold dry weight increase (P<0.001), respectively. Introducing collagen to agricultural soil increased root weight by 64.94% (P<0.01) (Fig. 3F). Additionally, agricultural soil amended with collagen displayed a 60.65% (P<0.05) greater increase in root weight than soils amended with chitin. Adding chitin to agricultural soil also yielded a root weight increase of 2.67% (P=0.86). In native soil, adding chitin yielded a 2-fold increase in root weight (P<0.001), while adding collagen yielded a 1.7-fold increase in root weight (P<0.01).

Under high inoculum pressure, agricultural soil also produced increased dry shoot weight, specifically by 1.1-fold (P<0.001) with chitin amendment and by 1.1-fold (P<0.001) with collagen (Fig. 1G). In native soil, the chitin treatment produced a 78% increase (P<0.01), while collagen produced a 65.5% increase (P<0.01) compared to the unamended, infected treatment. The roots in agricultural soil amended with collagen exhibited a 1.1-fold increase, while those with a chitin amendment showed a 1.4-fold increase (Fig. 1H). In native soils, chitin produced a 1.5-fold increase and collagen a 1.2-fold increase in root biomass.

The uptick in biomass and photosynthetic pigments, alongside a reduction in nematode eggs coincided with an alteration in the microbial community. In chitin-treated soils, there was a notable increase in the relative abundance of phyla like *Actinobacteria* (Fig 2A).

**Figure 2.**
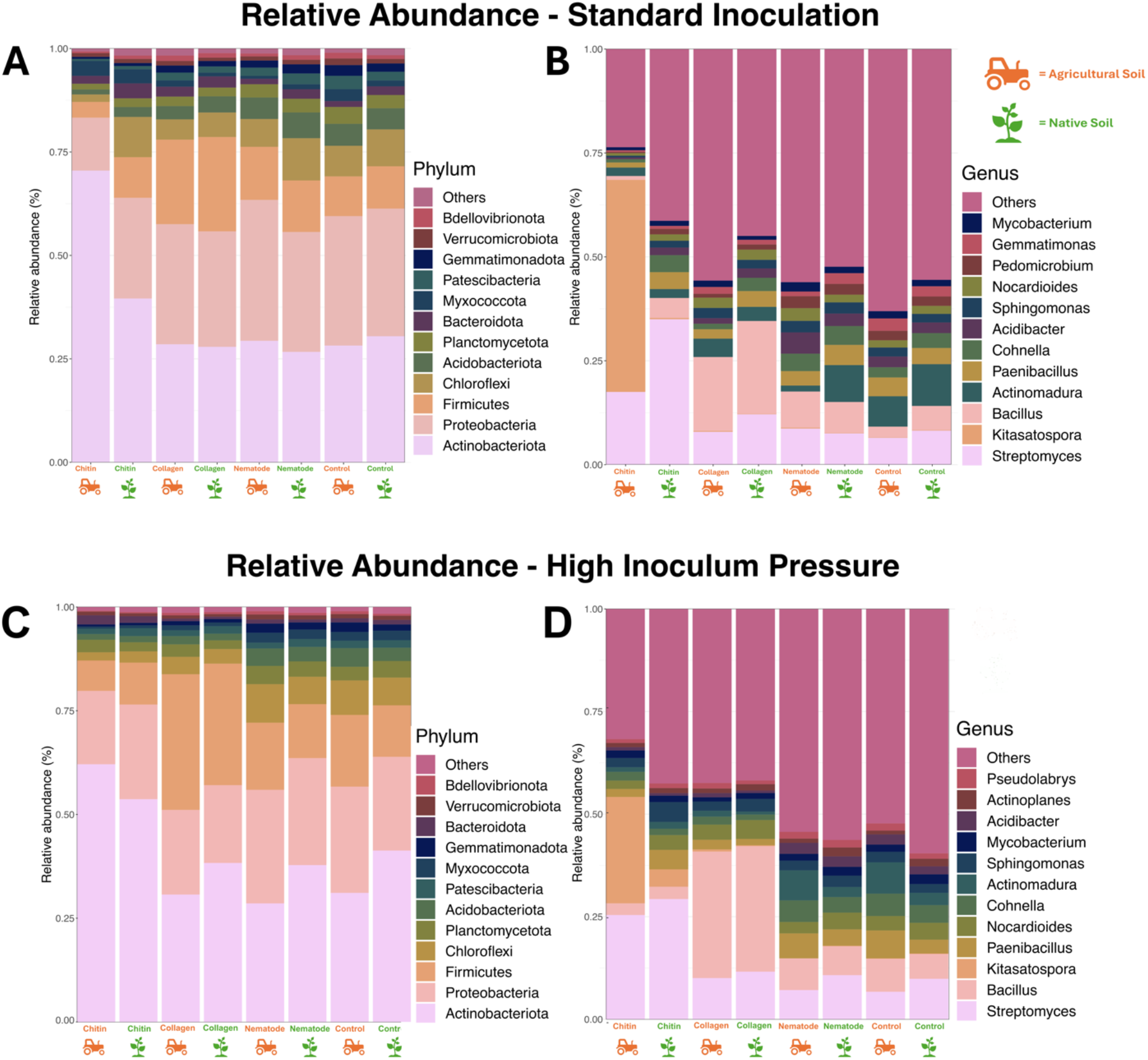
Relative abundance of operational taxonomic units (OTUs) of the twelve most abundant bacteria at the (A, C) phylum and (B, D) genus level. Bacterial community samples taken from tomato cv. Little Napoli rhizosphere at time of harvest. Comparison of standard inoculation (5,000 eggs) (A, B) and high inoculum pressure (50,000 eggs) (C, D). Means of 5 replicates.

Moreover, the genus *Kitasatospora* has particularly high abundance in agricultural soil treated with chitin, whereas native soil treated with chitin exhibits an increase in *Streptomyces* compared to other treatment groups (Fig. 2B). At the genus level, the collagen-treated group in both soil types displayed a pronounced increase in the genus *Bacillus* compared to other treatments. Similar results were observed in the experiment under high inoculum pressure (Fig. 2CD). In the agricultural soil treated with chitin, *Kitasatospora* dominated, whereas the collagen treatment resulted in a similar increase in *Bacillus* population. However, the *Bacillus* population was higher in the high-pressure experiment compared to the standard.

ITS sequencing revealed shifts in the abundance of fungi in response to experimental treatment groups. Considering phylum distribution, the standard inoculation collagen-treated groups exhibited an elevation in *Anthophyta* for both soil types (Fig. 3A).

**Figure 3.**
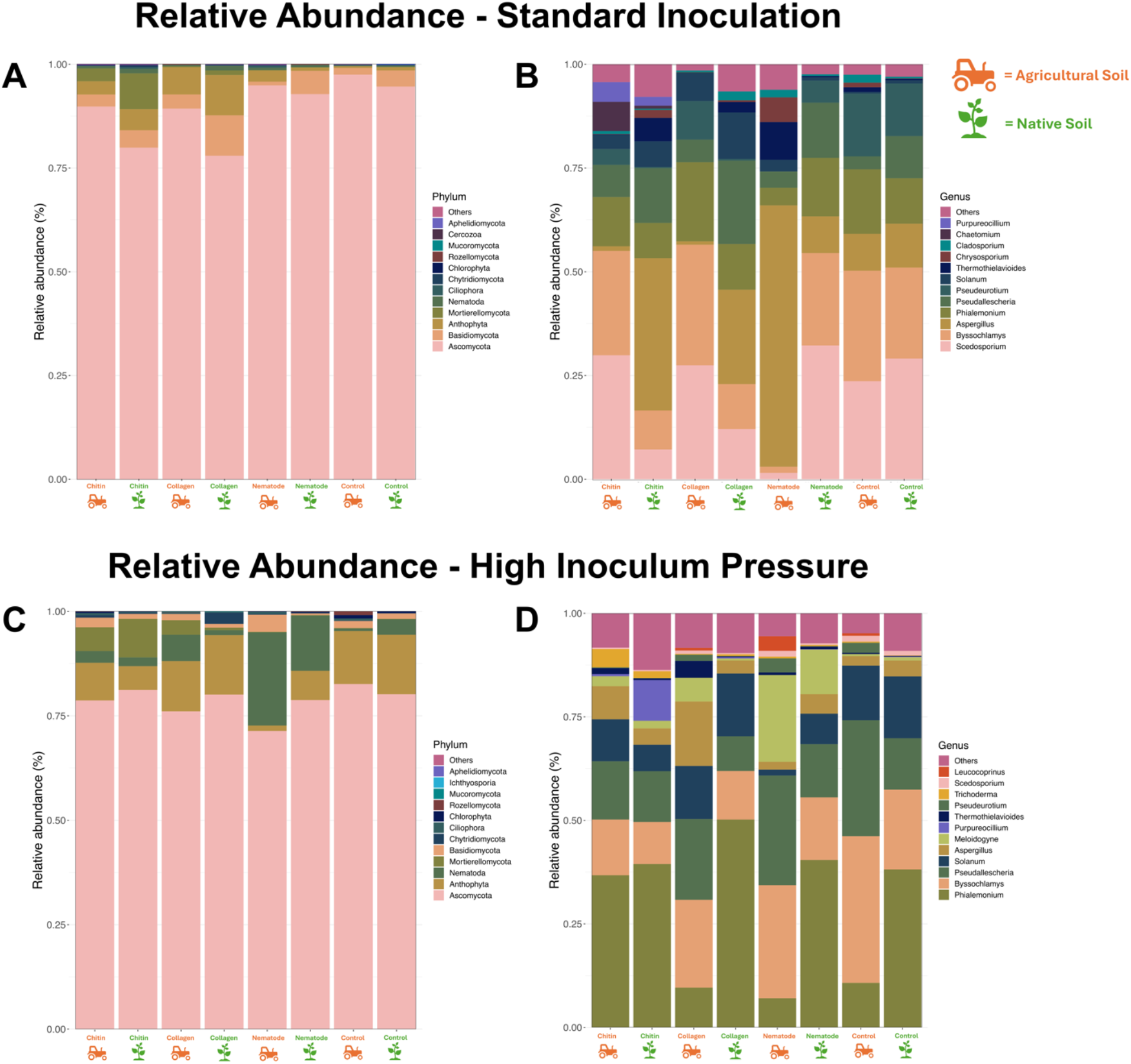
Relative abundance exhibiting the 12 most abundant fungal microbial assemblages at the phylum (A, C) and genus (B, D) level. Fungal community samples taken from tomato cv. Little Napoli rhizosphere at time of harvest. Comparison of standard inoculation (5,000 eggs) (A, B) and high inoculum pressure (50,000 eggs) (C, D). Means of 5 replicates.

In contrast, the chitin-treated groups showed a rise in *Mortierellomycota* compared to other groups. While native soil generally presented a higher abundance of *Basidiomycota*, the collagen-treated group in native soil surpassed this, boasting the highest abundance. Differences in genus were observed between amended soil types, with amended agricultural soil containing higher levels of *Byssochlamys* and *Scedosporium* while the amended native soil showed higher abundance of *Aspergillus* and *Pseudoallescheria* (Fig. 3B). The group under high inoculum pressure showed a similar trend with more alignment under soil type than specific treatment (Fig. 3D). This group also showed an overall increase in *Anthophyta*, with the most abundant groups being the control and collagen-treated groups (Fig. 3C). An increase in the phylum *Nematoda* was also seen, but most dramatically in the infected control groups for both soil types, corresponding with an increase in *Meloidogyne*. In the high inoculum group, the chitin amendment led to an increase in the *Phialemonium* spp. compared to the other treatments in the agricultural soil (Fig. 3D).

Along with these shifts in the amended soils, there was a decrease in diversity compared to the unamended groups, with chitin showing the lowest richness (Fig. 4A).

**Figure 4.**
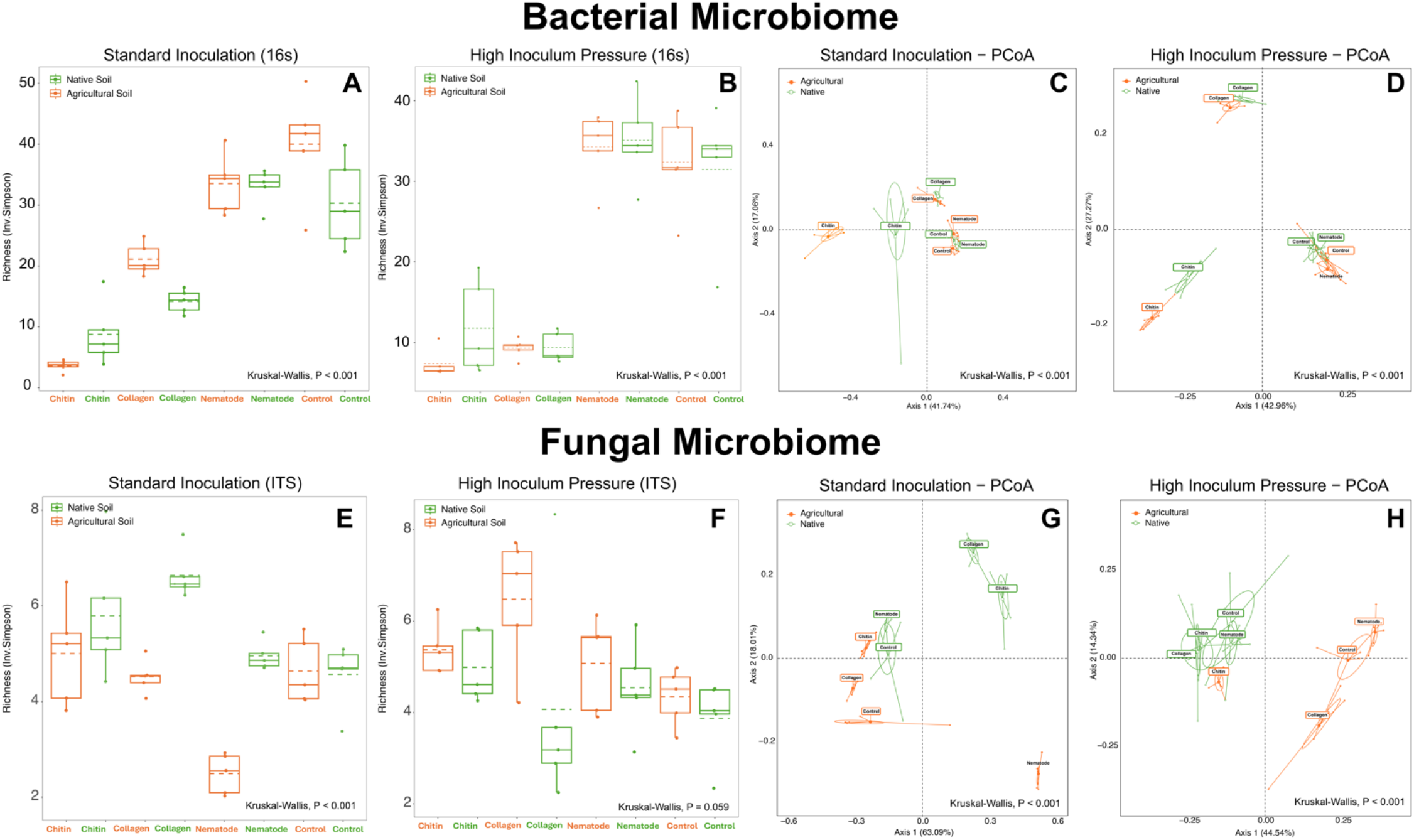
Visualization of microorganism diversity tomato cv. Little Napoli rhizosphere. Bacterial (A, B) and fungal (E, F) diversity under (A, E) standard (5,000 eggs) and (B, F) high inoculum pressure (50,000 eggs) through Inverse Simpson Plot. Principal coordinates analysis (PCoA) based on Bray-Curtis distance metric of bacterial (C, D) and fungal (G, H) diversity under (C, G) standard and (D, H) high inoculum pressure. The percentage of the variation explained by the plotted principal coordinates is indicated on the axes. Color indicates the treatment group, as indicated in the key.

In the experiment under high inoculum pressure, a similar decrease in diversity was observed in the amended groups, however, a stronger decrease was observed in the collagen-amended groups (Fig. 4B). No clear distinction was observed in the ITS data.

The microbiome profiles of the treatment groups showed statistically significant separation in diversity (permutational multivariate ANOVA [PERMANOVA], *P*=0.001; Fig. 4CDGH). In the bacterial microbiome, the amended groups demonstrated clear differentiation from each other and from the non-amended groups. The pronounced separation between treatment groups is primarily attributed to the highest Principal Coordinate 1 (PC1 = 47.74%). Under high inoculum pressure, the differentiation between groups is more distinct, with the collagen and chitin groups isolated from the unamended groups, attributed to the highest Principal Coordinate 1 (PC1 = 42.96%). In the fungal microbiome, a separation was not seen based on the treatment type, but instead by soil type.

The bacterial OTU-OTU networks revealed changes in the bacterial microbiome in response to soil type, inoculation intensity, and soil amendment. In the native soil under standard inoculation, amendments in the soil increased organisms such as *Streptomyces* Sd-13 and *Bacillus* spp. (Fig. 9GH).

A similar increase in *Bacillus* and *Streptomyces* groups were observed in the agricultural soil; however, in the group treated with chitin amendment, there was a notable shift towards *Kitasatospora* sp. 2391 emerging as the predominant organism (Fig. 5D). Under high inoculum pressure, similar outcomes were observed, demonstrating an increase in various *Bacillus* spp. in soils amended with collagen (fig 10). Additionally, in the agricultural soil, a rise in *Kitasatospora* was also noted, mirroring the trend seen in the standard inoculation group (fig. 5L).

**Figure 5.**
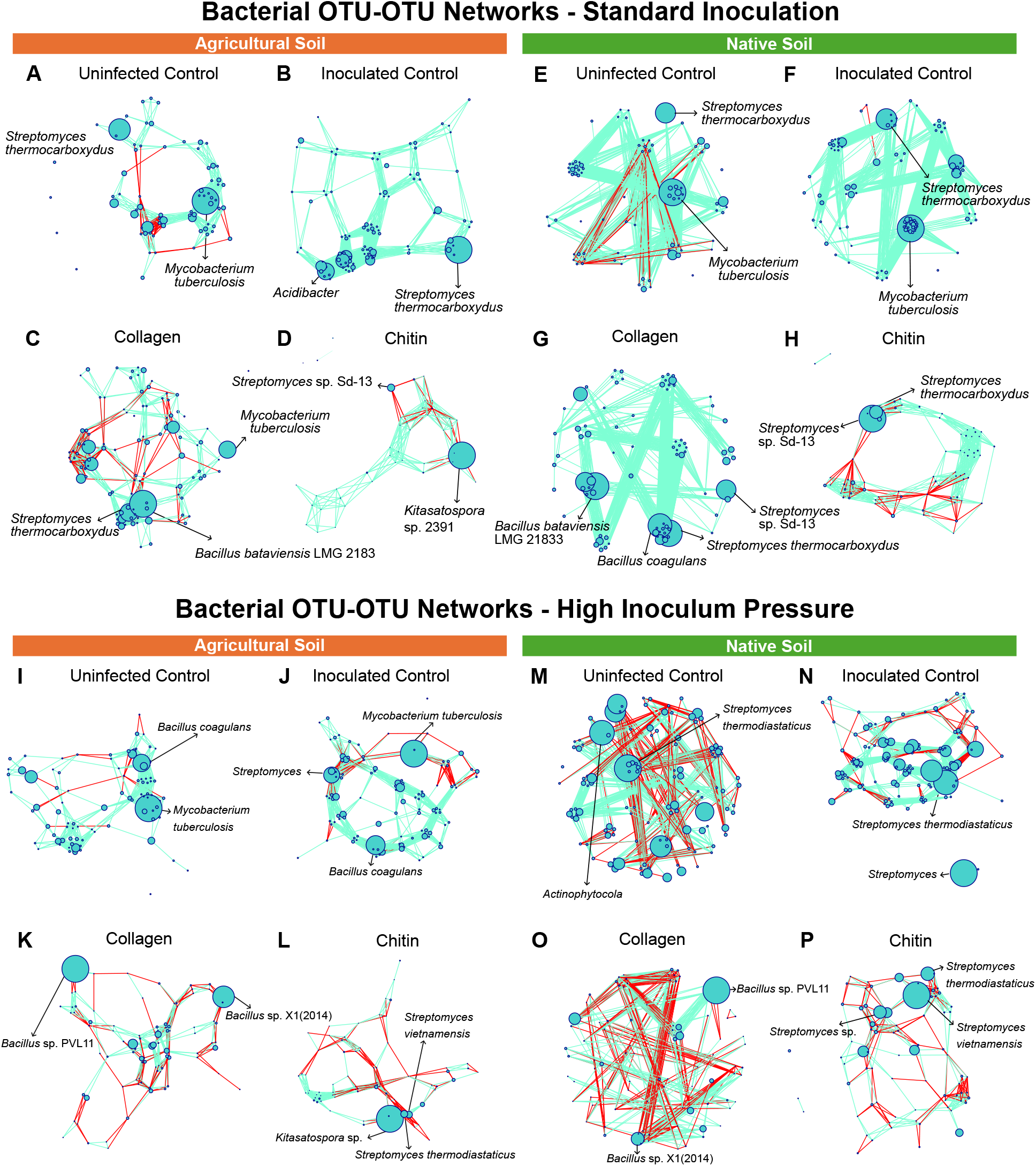
16S OTU-OTU network constructed from tomato (cv. Little Napoli) rhizosphere samples in agricultural (A-D, I-L) and native (E-H, M-P) soil under standard (5,000 eggs) (A-H) and high (50,000 eggs) (I-P) inoculation. Node size is proportional to relative abundance and edge color indicates positive (blue) and negative (red) correlations for the four treatments: negative control, inoculated control (Nematode), collagen-treated soil (Co) and chitin-treated soil (Ch) (five replicates per treatment).

In the fungal OTU-OTU networks, changes in relative abundance were less notable compared to the bacterial networks, as many genera remained consistent across different treatments (Fig. 6).

**Figure 6.**
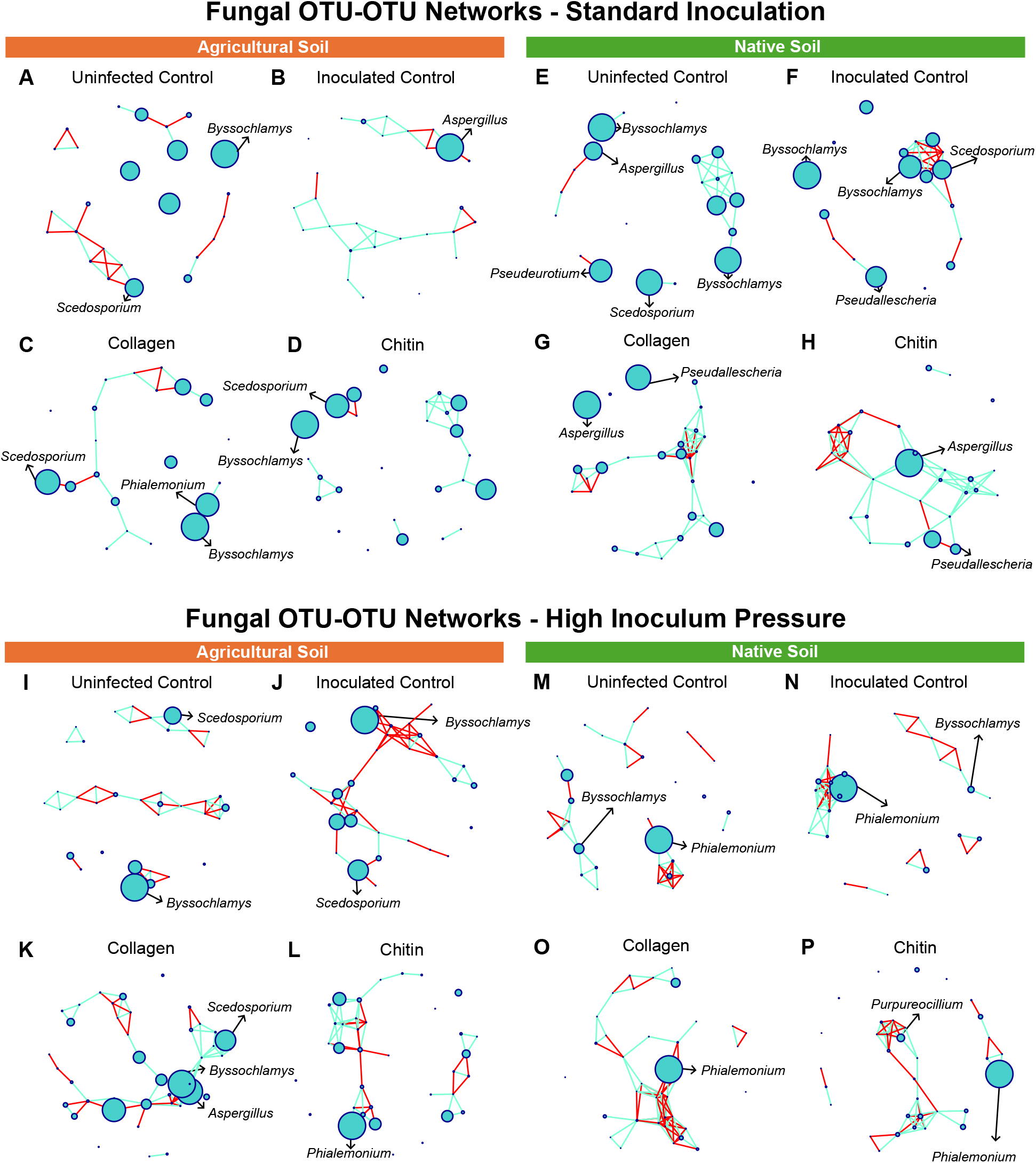
ITS OTU-OTU network constructed from tomato (cv. Little Napoli) rhizosphere samples in agricultural (A-D, I-L) and native (E-H, M-P) soil under standard (5,000 eggs) (A-H) and high (50,000 eggs) (I-P) inoculation. The node sizes correspond to the relative abundance and edge color corresponding to positive (blue) and negative (red) correlations for the four treatments: negative control, inoculated control (Nematode), collagen-treated soil (Co) and chitin-treated soil (Ch) (five replicates per treatment).

However, in the native soil under high inoculum pressure, *Phialemonium* emerged as the dominant organism (Fig. 6M-P). This genus also specifically dominated the chitin group in the agricultural soil under similar conditions (Fig. 6L).

Regarding overall network dynamics, more negative edges were observed under high inoculum pressure compared to standard inoculation scenarios, indicating the potential for more competitive interactions between microbes. (Table 3).

**Table 3:**
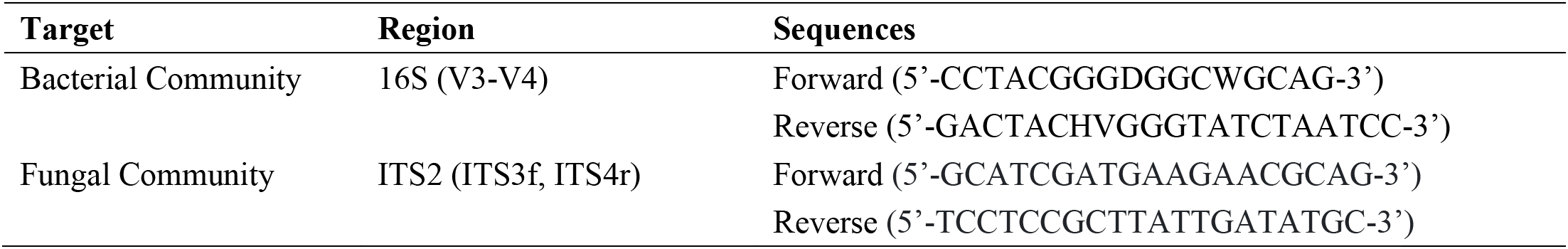
Target region to be used in amplicon sequencing for fungal and bacterial microbial communities.

**Table 3.** Summary of key network metrics for bacterial community analysis.

| Treatment | Positive Edges | Negative Edges | Total Connections | Average Node Degree | Number of Nodes |
| --- | --- | --- | --- | --- | --- |
| Standard Inoculation Agricultural Soil |  |  |  |  |  |
| Uninfected Control | 334 | 46 | 380 | 10.56 | 72 |
| Inoculated Control | 1098 | 0 | 1098 | 22.875 | 96 |
| Collagen | 421 | 124 | 545 | 10.8 | 101 |
| Chitin | 163 | 29 | 192 | 9.37 | 41 |
| Standard Inoculation Native Soil |  |  |  |  |  |
| Uninfected Control | 229 | 12 | 241 | 5.95 | 81 |
| Inoculated Control | 1339 | 2 | 1341 | 27.65 | 97 |
| Collagen | 877 | 0 | 877 | 21.13 | 83 |
| Chitin | 469 | 71 | 540 | 14.8 | 73 |
| High Inoculum Pressure Agricultural Soil |  |  |  |  |  |
| Uninfected Control | 492 | 17 | 509 | 11.31 | 90 |
| Inoculated Control | 627 | 93 | 720 | 13.46 | 107 |
| Collagen | 241 | 128 | 369 | 8.11 | 91 |
| Chitin | 235 | 112 | 347 | 8.9 | 78 |
| High Inoculum Pressure Native Soil |  |  |  |  |  |
| Uninfected Control | 616 | 292 | 908 | 14.76 | 123 |
| Inoculated Control | 962 | 108 | 1070 | 15.97 | 134 |
| Collagen | 350 | 214 | 564 | 12 | 94 |
| Chitin | 182 | 132 | 314 | 8.05 | 78 |

Key network metrics varied among treatment groups, highlighted in table 3, indicate distinct microbial community responses.

In the fungal microbial networks, the high inoculum pressure groups also resulted in more negative edges and a competitive microbial environment (Table 4).

**Table 4:** Gall incides assessed at harvest from the roots of tomato cv. Little Napoli plants under standard (5,000 eggs) and high inoculation pressure (50,000 eggs)

| Treatment | Native soil |  |  |  | Agricultural Soil |  |  |  |
| --- | --- | --- | --- | --- | --- | --- | --- | --- |
|  | Uninfected Control | Infected Control | Chitin | Collagen | Uninfected Control | Infected Control | Chitin | Collagen |
| Brito et al., 2020 |  |  |  |  |  |  |  |  |
| Standard Inoculation | 1.75 | 5 | 4.875 | 4.75 | 0 | 5 | 5 | 4.625 |
| High Inoculum Pressure | 1.2 | 4.8 | 4.6 | 4.8 | 1.6 | 5 | 5 | 5 |
| Hussey & Janssen., 2002 |  |  |  |  |  |  |  |  |
| Standard Inoculation | 0.625 | 4.375 | 3.375 | 2.625 | 0 | 3.75 | 3.125 | 2.375 |
| High Inoculum Pressure | 0.6 | 4.4 | 4.2 | 3.6 | 0.8 | 4.8 | 5 | 3.6 |

**Table 4.** Summary of key network metrics for fungal community analysis.

| Treatment | Positive Edges | Negative Edges | Total Connections | Average Node Degree | Number of Nodes |
| --- | --- | --- | --- | --- | --- |
| Standard Inoculation Agricultural Soil |  |  |  |  |  |
| Uninfected Control | 9 | 16 | 25 | 2 | 25 |
| Inoculated Control | 27 | 7 | 34 | 2.43 | 28 |
| Collagen | 13 | 5 | 18 | 1.71 | 21 |
| Chitin | 22 | 2 | 24 | 1.85 | 26 |
| Standard Inoculation Native Soil |  |  |  |  |  |
| Uninfected Control | 24 | 3 | 27 | 2.57 | 21 |
| Inoculated Control | 14 | 14 | 28 | 3.11 | 18 |
| Collagen | 45 | 12 | 57 | 3.93 | 29 |
| Chitin | 52 | 16 | 68 | 4.53 | 30 |
| High Inoculum Pressure Agricultural Soil |  |  |  |  |  |
| Uninfected Control | 33 | 23 | 56 | 2.95 | 38 |
| Inoculated Control | 20 | 35 | 55 | 3.44 | 32 |
| Collagen | 56 | 22 | 78 | 3.8 | 41 |
| Chitin | 39 | 20 | 59 | 3.28 | 36 |
| High Inoculum Pressure Native Soil |  |  |  |  |  |
| Uninfected Control | 24 | 20 | 44 | 2.67 | 33 |
| Inoculated Control | 36 | 30 | 66 | 4.26 | 31 |
| Collagen | 47 | 40 | 87 | 4.7 | 37 |
| Chitin | 33 | 24 | 57 | 3.35 | 34 |

The bacterial Phenotype-OTU network indicates OTUs significantly (P<0.05) associated with lower disease severity (Fig. 9AD)

Two OTU’s, *Paradeviosa shaoguanesis* and *Streptomyces* sp. were identified as both associated with lower disease incidence and a part of the overall core microbiome. A phenotype-OTU analysis was performed on the ITS data, revealing 10 significant associations (P < 0.05) with low disease incidence (Fig. 7D). These organisms encompassed four different phyla, including three unidentified fungi. *Mortierellomycotina* was found in the highest abundance, while the strongest coefficient was attributed to *Chytridiomycota*.

**Figure 7.**
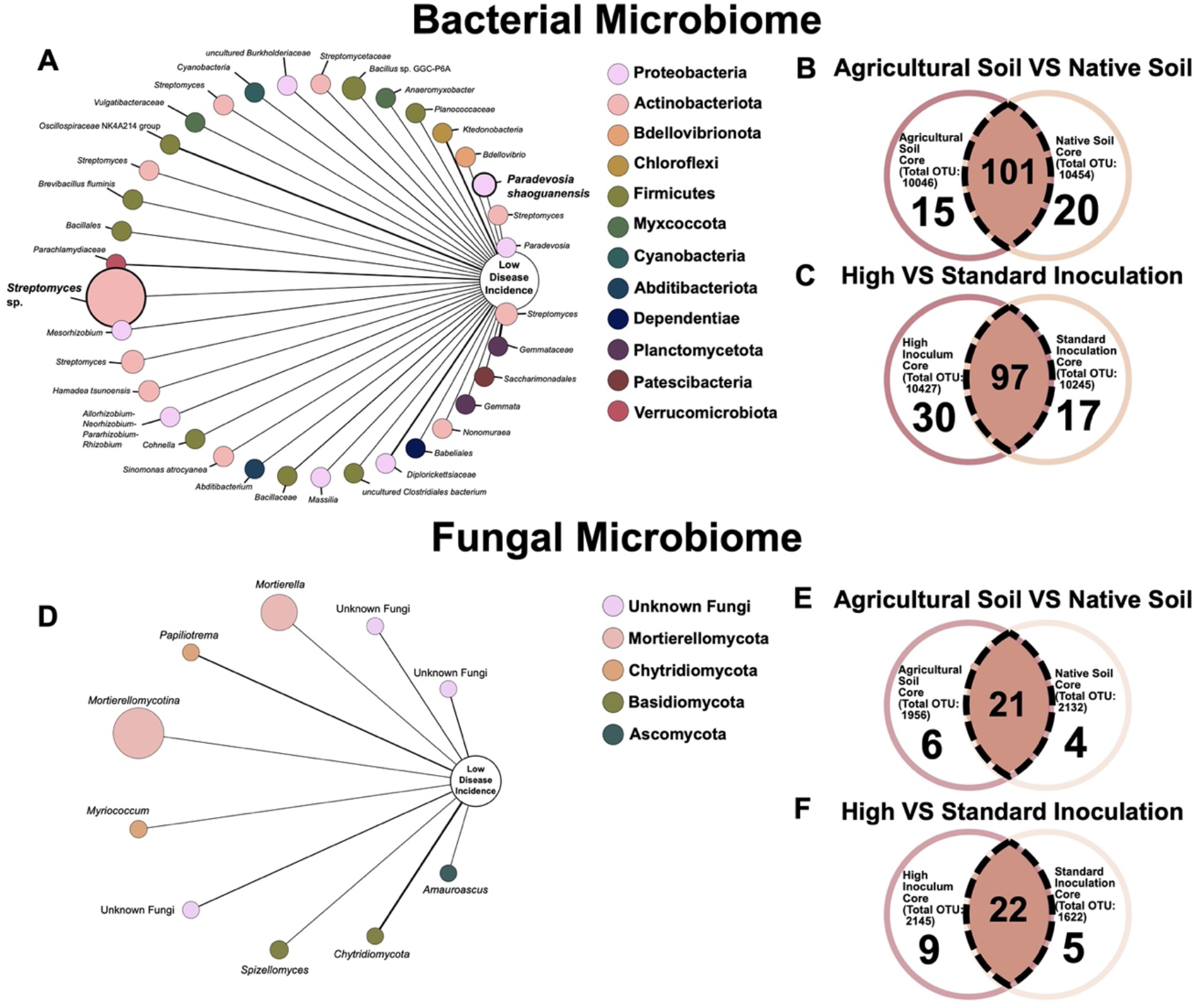
The OTU-phenotype network uncovers (A) 36 and (B) 10 significant OTUs (Operational Taxonomic Units) (*P*<0.05) strongly associated with the log odds probability, as determined by logistic regression analysis, of decreased disease resistance in plants infected with *Meloidogyne enterolobii*. This association was established by considering treatments falling within the bottom quartile of eggs per root gram. Nodes in the network are color-coded according to the corresponding phylum of the OTU, while node size reflects its relative abundance. The width of edges signifies the strength of the probability coefficient, with solid lines indicating positive associations. Additionally, a bolded font and node indicate members of the core microbiome. Bacterial (B, C) and fungal (E, F) Venn diagram displaying number of OTUs of soil types (B, E) and experiment (C, F) found and the core microbiome between them at a 0.001 relative abundance detection limit and a 50% prevalence.

A core microbiome across soil types was also identified, comprising 101 OTUs in the 16s data and 21 in the ITS data at a 0.001 abundance detection limit and 50% prevalence (Fig. 7BE). Between experiments, 97 bacterial OTUs and 22 fungal OTUs were found to compose the core microbiome shared between standard inoculation and high inoculum pressure under the same parameters (Fig. 7CF).

Untargeted soil metabolomic analysis, a comprehensive approach used to identify and quantify a wide range of metabolites present in soil samples without prior knowledge of what these metabolites might be, was conducted, revealing a notable disparity between high inoculum pressure and standard inoculation. The high inoculum pressure group exhibited lower expression of metabolic groups (classes) whereas the standard inoculation group showed increased expression (Fig. 10AC).

Additionally, in the PCoA analysis of metabolites, distinct separation between the experiments was observed. The pronounced segregation between treatment groups is primarily attributed to the highest Principal Coordinate 1 (PC1 = 26.95%) (Fig. 8B). The most prominent metabolic group was amino acids and their metabolites, likely indicating higher activity levels in the soil.

**Figure 8.**
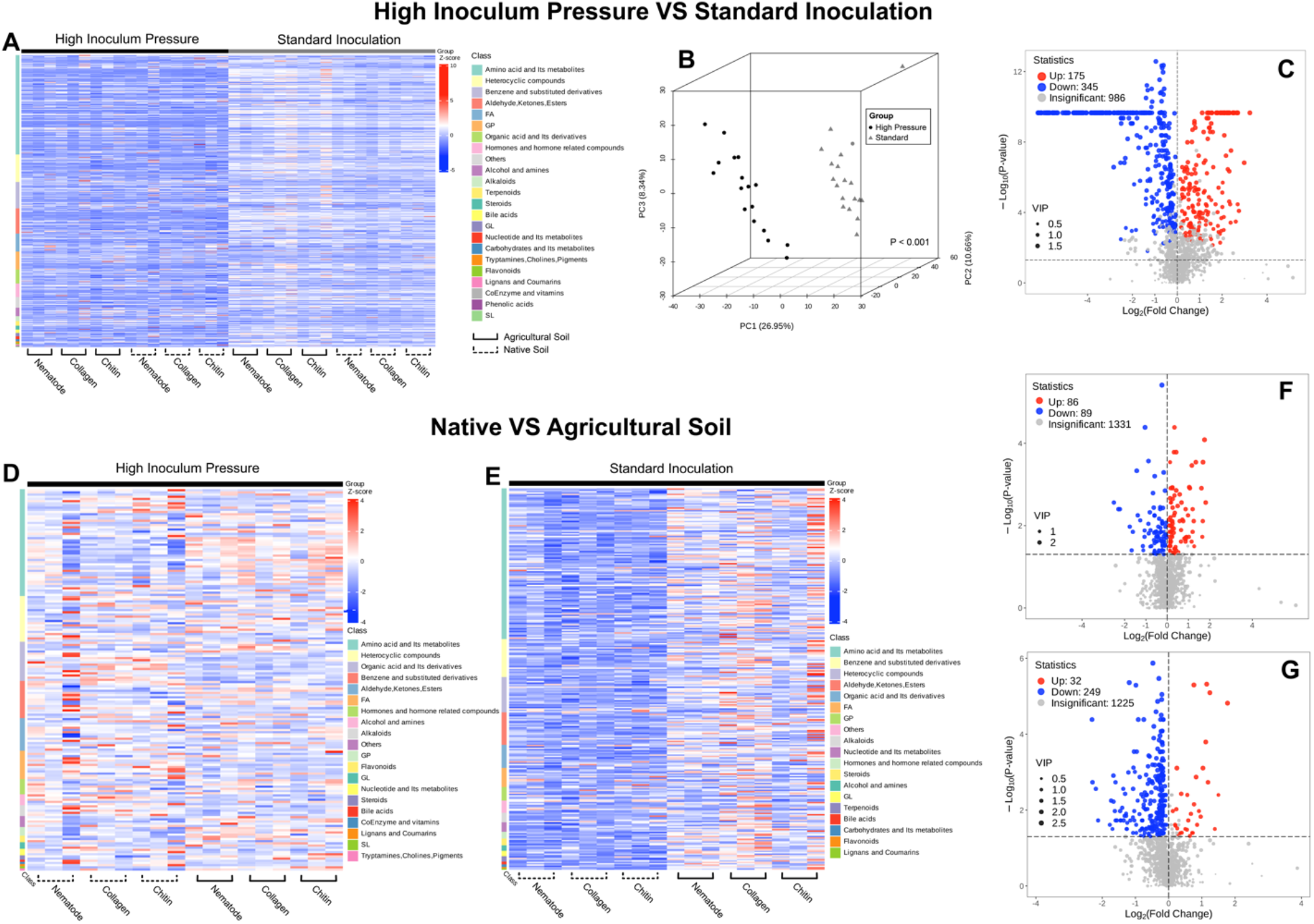
Untargeted soil metabolomics: differential analysis between high inoculum pressure and standard inoculation conditions This figure contains comparisons between (ABC) two nematode inoculation levels and (DEFG) two soil types. The analyses include: (A) A heatmap displaying the expression levels of various metabolic groups. (B) A principal coordinates analysis (PCoA) illustrating the clustering of metabolic profiles. (C) A volcano plot that identifies significantly upregulated and downregulated metabolites. (D) A heatmap displaying the expression levels of various metabolic groups under high inoculum pressure (50,000 eggs). (E) A heatmap displaying the expression levels of various metabolic groups under standard inoculation (5,000 eggs. (F) A volcano plot identifying significantly upregulated and downregulated metabolites under high inoculum pressure. (G) A volcano plot identifying significantly upregulated and downregulated metabolites under standard inoculation.

In comparing untargeted metabolomics under standard inoculation across soil types, a visible difference was evident, with decreased expression observed in the native soil, contrasting with elevated expression in agricultural soil (Fig. 8E). However, this contrast was not evident under conditions of high inoculum pressure. While metabolite expression remained similar in the agricultural soil under both conditions, the native soil exhibited heightened expression levels under high inoculum pressure as well (Fig. 8D). This trend was further illustrated in the volcano plots, where numerous metabolites were down-expressed in the standard inoculation group (Fig. 8G), while the high inoculum pressure group showed a more balanced distribution of up- and down-expressed metabolites (Fig. 8F). Amino acids were also the most commonly observed metabolic group; however, in the high inoculum pressure treatment, amino acids were observed to be a smaller majority, leading to a more even metabolic fingerprint.

## Discussion

In this study we observed reduced nematode egg counts with the addition of the amendments, collagen and chitin, for both soil types and for both levels of inoculum pressure.

This highlights the potential of soil amendments to alter the soil microbiome and suppress *M. enterolobii* populations. An increase in *Meloidogyne* spp. inoculum is directly proportional to yield loss in a tomato plant (Kankam & Adomako, 2014), thus the ability to decrease inoculum load shows promise for future methods of biocontrol. Under standard inoculation conditions, collagen reduced nematode eggs per gram of root in agricultural soil, demonstrating its strong evidence for efficacy in nematode management, supporting results found by Galper et al. (1990). There was weaker evidence for the effect of chitin in agricultural soil (P=0.67), but a consistent downward trend was noted, suggesting a potential suppressive effect that warrants further exploration. Moreover, we found that native and agricultural soils respond differently to the soil amendment. Both amendments exhibited strong evidence of effectiveness in native soil under standard inoculation. According to a study by Chialva et al. (2018), the microbiome of native soils can induce a “state of alert” in tomatoes, enhancing responses to soilborne pathogens.

Under high inoculum pressure, we observed a substantial decrease in eggs for both amendments in agricultural soil, affirming their vigor against elevated disease challenges. Although reductions in native soil under these high-pressure conditions did not achieve statistical significance, the consistent trend of decreases across various conditions indicates a continued potential for nematode suppression. These findings indicate that the efficacy of collagen and chitin in reducing nematode populations may vary depending on soil type, and therefore different soil microbiome, suggesting the indirect effect of the amendments in controlling the plant-parasitic nematode. Prior research has suggested that alterations in soil cultivation practices can impact soil suppressiveness and alter the microbial community to favor defense against *M. enterolobii* (Pasche et al., 2023). Thus, the soil microbes present in different soil types may differ in their use of collagen and chitin, impacting their efficacy against *M. enterolobii* infection.

Though we found evidence that suggests a microbial component in M. enterolobii control, it is important to also consider the role of chitin-induced resistance (CIR). Chitin, upon application, may have been recognized by the plant’s pattern recognition receptors (PRRs), which can initiate defense mechanisms against nematodes (Riseh et al., 2024). Interestingly, the systemic acquired resistance (SAR) pathway has been shown to be less effective against root-knot nematodes due to the nematode’s ability to suppress the expression of key PR genes (Sanz-Alférez et al., 2007). However, due to the variability of CIR depending on host and pathogen species, different or additional defense pathways may be activated to provide a response against nematode infections (Riseh et al., 2024). The dual impact of beneficial microorganisms and CIR could synergistically contribute to the observed reduction in nematode egg counts. Further research into the CIR mechanism and its effectiveness against *M. enterolobii* in tomato could provide valuable insights into chitin not only enhancing microbial activity in the soil, but also directly triggering plant immune responses, providing a comprehensive approach to nematode management.

There was strong evidence for increased plant biomass under both standard inoculation and high inoculum pressure conditions when soil was amended with collagen and chitin. These results suggest that the amendments alleviate plant stress through disease mitigation and potential biostimulatory effects. Chitin is known to enhance plant vigor and tolerance to stress, as documented by Pichyangkura & Chadchawan (2015) and Shahrajabian et al. (2021). Similarly, collagen promotes plant growth when broken down into bioavailable forms by soil microorganisms, functioning as a biostimulant (Ambrosini et al., 2021). Increases in chlorophyll content noted in amended soils likely result from these biostimulatory effects. Liopa-Tsakalidi et al. (2010) demonstrated that soil supplementation with chitin could elevate chlorophyll levels by up to 60%, which correlates with enhanced photosynthetic capacity and potentially improved plant yield. Likewise, collagen has been shown to boost chlorophyll production, thereby facilitating increased photosynthetic activity and overall plant health (Ambrosini et al., 2021).

The additions of collagen and chitin, resulting in diminished microbial diversity and richness under both standard and high inoculum pressure conditions, signify a shift in the soil microbiome towards an increase in high-abundance taxa compared to a more balanced distribution (Figure 7). The microbial community differentiation between amended and unamended groups is also highlighted in the principal coordinates analysis. This differentiation is primarily driven by shifts along Principal Coordinate 1, which captures the largest portion of the variance, indicating a strong influence on community composition driven by the amendments.

The shift in microbial communities appears effective, particularly under high inoculum pressures, further suggesting that the amendments might be promoting microbial taxa that are suppressive to nematodes or otherwise beneficial for plant health.

In groups treated with chitin, a noticeable shift in the microbiome occurred, possibly due to the rise in chitin-degrading enzymes. In the standard inoculation group, there was an increase in *Streptomyces* when chitin was introduced into the soil. Moreover, under high inoculum pressure, this trend persisted, with *Streptomyces* experiencing even more substantial growth.

Most streptomycetes are known to secrete chitinases (Schrempf, 2001), and many *Streptomyces* strains have been specifically identified for their production of chitin-degrading enzymes (Kumar et al., 2020). *Streptomyces* was the predominant genus in the native soil group; however, in agricultural soil, *Kitasatospora* dominated. Sawaguchi et al. (2015) found that both *Streptomyces* and *Kitasatospora* exhibited strong increases with the addition of chitosan to the soil. Moreover, most chitosan-degrading bacteria isolated were identified as *Streptomyces*, strongly suggesting the involvement of these two groups in chitosan degradation. Specific strains of *Kitasatospora*, such as *Kitasatospora setae*, have been identified as producers of chitinases (Zitouni et al., 2017).

Hui et al. (2020) identified *Bacillus* and *Kitasatospora* as opportunistic species in chitin degradation. *Bacillus* also increased in the collagen-amended soil, and many strains of *Bacillus* are known to produce collagenases (Evans and Wardlaw, 1953). Many collagenolytic *Bacillus* strains have been identified (Hoppe et al., 2021; Mohindra et al., 2020, Tran & Nagano, 2002). This increase in *Bacillus* when exposed to a collagen soil amendment may indicate an increase in collagenolytic *Bacillus* species. The increase of *Phialemonium* genera shown in the abundance and OTU-OTU networks (Fig. 3D and Fig. 6L-P) may related to the ability of the *Phialemonium* to antagonize root-knot nematodes, as shown in previous research (Zhou et al. 2018).

The network metrics in Table 3 indicate that the microbial communities in amended soils are influenced by these collagen and chitin treatments. Previous studies have highlighted the shift in microbial community dynamics when exposed to soil amendments (Zhou et al., 2019; Francioli et al., 2016; Hallmann et al., 1999). In the study conducted by Zhou et al. (2019), the group found that addition of the biochar amendment increased microbial network complexity. In our study, under standard inoculation, the result mirrored findings by Zhou et al. (2019), as collagen amendment in agricultural soil increased the number of both positive and negative edges. Conversely, the chitin amendment produced fewer overall connections, but the presence of negative edges in the OTU-OTU networks in the chitin-amended group may reflect selection for nematode-suppressive microbes. In native soils under the same conditions, collagen treatment results in a high number of positive edges and an absence of negative edges, could be indicative of a network of mutualistic interactions. In native soils under the same conditions, collagen treatment results in a high number of positive edges and an absence of negative edges, indicative of a network of mutualistic interactions. Chitin introduces a notable number of negative edges, which may imply a shift towards a more competitive microbial environment, possibly favoring nematode antagonistic taxa. However, Poudel et al. (2016) explain that observed network structures in microbiome models can reflect a combination of biological interactions and shared environmental preferences, rather than direct mutualistic relationships. Thus, while the positive edges in collagen-treated soils suggest potential mutualistic interactions, they may also indicate taxa that thrive in similar environmental niches. In addition to showing a change in the soil microbial composition with the importation of soil amendments, we also saw a change in the soil microbial function with the metabolites assessed. Overall, these findings suggest strong evidence that the application of collagen and chitin amendments influences both the structure and function of soil microbial communities.

Under high inoculum pressure, the number of negative edges increased with collagen treatment in the agricultural soil, which could reflect an adaptive response of the microbial community under biotic stress, potentially directed towards inhibiting nematode infection.

Chitin’s impact appears similar, with an increase in negative interactions that could influence microbial competition. In native soil experiencing high inoculum pressure, both collagen and chitin treatments have fewer total connections and lower average degree, suggesting the establishment of a more specialized microbial network.

The bacterial phenotype-OTU network displays 36 OTUs (P<0.05) associated with lower disease severity, highlighting the association between specific bacterial taxa and decreased disease incidence (Fig. 9AD). Notably, two OTUs, *Paradeviosa shaoguanesis* and *Streptomyces sp.*, were both associated with lower disease incidence and identified as members of the core microbiome. This dual role underscores their potential importance in soil health and disease suppression. The identification of a larger number of significant bacterial OTUs compared to fungal OTUs can be attributed to several factors. Bacterial communities generally exhibit greater diversity and abundance, which might enhance the likelihood of detecting significant associations with disease resistance. Additionally, bacteria may respond more sensitively to inoculum pressure, or methodological biases might favor the detection and analysis of bacterial DNA over fungal DNA.

Conversely, the phenotype-OTU analysis for fungal communities revealed only 10 associations (P<0.05) with low disease incidence (Fig. 7D). These fungal OTUs encompassed four different phyla, including three unidentified fungi. The phylum *Mortierellomycotina* was found in the highest abundance, whereas the strongest coefficient was attributed to *Chytridiomycota*. This lower number of significant fungal OTUs might be due to various factors such as lower diversity or methodological challenges in detecting and characterizing fungi in the microbiome.

The core microbiome consists of groups of microorganisms that establish essential interactions, which can be leveraged to enhance microbial functions (Toju et al., 2018). The identification of core microbiomes in agricultural soils, comprising both bacterial and fungal constituents, offers promising avenues for enhancing plant health and disease management strategies, such as use in potential synthetic microbial communities (Wang et al., 2023). The core microbes are favorable for such management strategies due to their strong interactions with plant functions and can aid in better recruitment and establishment of other beneficial microbes (Toju et al., 2018). Among the 10,817 OTUs, this analysis identified 97 bacterial and 22 fungal OTUs as part of the “core microbiome” across different experimental conditions— under both standard inoculation and high inoculum pressure scenarios as depicted in Figure 7 (C and F). The same analysis was conducted between soil types revealing core bacterial (101 OTUs) and fungal (21 OTUs) communities across agricultural and native soil types (Figure 7BE). This conserved microbial cohort suggests a fundamental relationship between these taxa and plant health, irrespective of the level of disease pressure and soil type.

Untargeted soil metabolomic analysis has highlighted disparities between conditions of high inoculum pressure and standard inoculation. Under high inoculum pressure, there is a marked decrease in the expression of metabolic groups compared to standard inoculation conditions. This reduction may indicate a suppression or reallocation of metabolic functions, possibly as a response to stress or disease conditions in the soil environment (Withers et al., 2020). The pronounced separation observed in the Principal Coordinates Analysis (PCoA) underscores that the metabolic responses to high inoculum pressure differ significantly from those under standard conditions (Fig. 8B). This clustering could reflect adaptive or defensive modifications in the soil microbiome, which are vital for understanding soil resilience and health under stress (Withers et al., 2020).

Untargeted metabolomic analysis also gave substantial insights into the differential metabolic responses of native and agricultural soils under varying disease pressure. Amino acids and their metabolites were the predominant class identified in the metabolomic results, with this class being even more pronounced under standard inoculation conditions. An increase in amino acid concentration in the soil can be linked to soil microbial activity and nutrient uptake (Vinolas et al., 2001; Sauheitl et al., 2009). In standard inoculation conditions, the native soil exhibited reduced metabolic activity. This difference may be due to the soil having a lower baseline metabolic activity due to inherent limitations in nutrient availability or microbial diversity due to less anthropogenic influence (Pasche et al., 2023). However, under high inoculum pressure, native soil demonstrated a notable increase in metabolic activity, suggesting a possible adaptive response to stress that aligns its metabolic profile more closely with that of agricultural soil.

This study represents the first comprehensive analysis of the response of soil microbiome (bacteria and fungi) structure and function to chitin and collagen amendments under different soil conditions and inoculum pressure using *Meloidogyne enterolobii* as a model soilborne parasite.

The results of this study demonstrate the potential of collagen and chitin amendments for mitigating nematode infection and promoting overall plant health. Both amendments displayed efficacy in reducing nematode egg counts, with varying efficacy observed in each soil type, which highlights the importance of soil type in microbiome response to amendments. The observed shifts in the soil microbiome suggest that these amendments may indirectly suppress nematodes by favoring the growth of specific microbial taxa. The identification of a core microbiome shared across treatments and soil types offers valuable insights for developing novel biocontrol agents against *M. enterolobii*. Future research should investigate the function of specific microbials in suppressing the nematode. Network analysis can help to identify potentially synergistic combinations of taxa (Poudel et al 2016, Poudel et al 2023), since a potentially more effective strategy would be using a combination of isolates as a synthetic microbial community (SynComs) with complementary functions (Martins et al. 2023), Considering both bacteria and fungi in SynComs has been shown to provide better results (Zhou et al. 2022).

## Acknowledgements

This work was supported by the USDA National Institute of Food and Agriculture (NIFA) project no. 2021-68013-33758 and hatch project no. 1024881. The authors are also thankful to Dr. Janete Brito for providing the nematode inoculum, Dr. Gary Vallad for giving us access to the fields for soil collection, and Mason Trub for assisting with a part of the greenhouse assay.

## REFERENCES

1. Ambrosini, S., Sega, D., Santi, C., Zamboni, A., Varanini, Z., & Pandolfini, T. (2021). Evaluation of the potential use of a collagen-based protein hydrolysate as a plant multi-stress protectant. Frontiers in plant science, 12, 63.

2. Brito, J. A., Desaeger, J., & Dickson, D. W. (2020). Reproduction of on selected root-knot nematode resistant sweetpotato () cultivars. Journal of Nematology, 52(1), 1–6.

3. Bui, H. X., & Desaeger, J. A. (2021). Host suitability of summer cover crops to Meloidogyne arenaria, M. enterolobii, M. incognita and M. javanica. Nematology, 24(2), 171–179.

4. Cetintas, R., Kaur, R., Brito, J. A., Mendes, M. L., Nyczepir, A. P., & Dickson, D. W. (2007). Pathogenicity and reproductive potential of Meloidogyne mayaguensis and M. floridensis compared with three common Meloidogyne spp. Nematropica, 21–32.

5. Chialva, M., Salvioli di Fossalunga, A., Daghino, S., Ghignone, S., Bagnaresi, P., Chiapello, M., … & Bonfante, P. (2018). Native soils with their microbiotas elicit a state of alert in tomato plants. New Phytologist, 220(4), 1296–1308.

6. Csardi G, Nepusz T (2006). “The igraph software package for complex network research.” InterJournal, Complex Systems, 1695. https://igraph.org.

7. Ekanayake, H. M. R. K., & Di Vito, M. (1984). Effect of population densities of Meloidogyne incognita on growth of susceptible and resistant totamo plants. Nematologia mediterranea.

8. Evans, D. G., & Wardlaw, A. C. (1953). Gelatinase and collagenase production by certain species of Bacillus. Microbiology, 8(3), 481–487.

9. Francioli, D., Schulz, E., Lentendu, G., Wubet, T., Buscot, F., & Reitz, T. (2016). Mineral vs. organic amendments: microbial community structure, activity and abundance of agriculturally relevant microbes are driven by long-term fertilization strategies. Frontiers in microbiology, 7, 215318.

10. Galper, S., Cohn, E., Spiegel, ∼ Y, & Chet, I. (1991). A Collagenolytic Fungus, Cunninghamella elegans, for Biological Control of Plant-parasitic Nematodes I. In JOURNAL OF NEMATOLOGY (Vol. 23, Issue 3).

11. Galper, S., Cohn, E., Spiegel, Y., & Chet, I. (1990). Nematicidal effect of collagen-amended soil and the influence of proteasëand collagenase (l). Revue Néwzatol, 13(1), 67–71.

12. Hallmann, J., Rodrıguez-Kábana, R., & Kloepper, J. W. (1999). Chitin-mediated changes in bacterial communities of the soil, rhizosphere and within roots of cotton in relation to nematode control. Soil Biology and Biochemistry, 31(4), 551–560.

13. Hoppe, I. J., Brandstetter, H., & Schönauer, E. (2021). Biochemical characterisation of a collagenase from Bacillus cereus strain Q1. Scientific Reports, 11(1), 4187.

14. Hui, C., Jiang, H., Liu, B., Wei, R., Zhang, Y., Zhang, Q., … & Zhao, Y. (2020). Chitin degradation and the temporary response of bacterial chitinolytic communities to chitin amendment in soil under different fertilization regimes. Science of the total environment, 705, 136003.

15. Hussey R. S., Barker K. R. (1973) A comparison of methods of collecting inocula of Meloidogyne spp., including a new technique. Plant Disease Reporter. 57:1025–1028.

16. Hussey, R. S., & Janssen, G. J. W. (2002). Root-knot nematodes: Meloidogyne species. In Plant resistance to parasitic nematodes (pp. 43–70). Wallingford UK: Cabi Publishing.

17. Kankam, F., & Adomako, J. (2014). Influence of inoculum levels of root knot nematodes (Meloidogyne spp.) on tomato (Solanum lycopersicum L.). Asian Journal of Agriculture and Food Sciences, 2(2).

18. Kumar, M., Kumar, P., Das, P., Solanki, R., & Kapur, M. K. (2020). Potential applications of extracellular enzymes from Streptomyces spp. in various industries. Archives of Microbiology, 202, 1597–1615.

19. Lahti, L., & Shetty, S. (2017). microbiome R package. Bioconductor. 10.18129/B9.bioc.microbiome

20. Lichtenthaler, H., & Wellburn, A. R. (1984). Formulae and program to determine total carotenoids and chlorophylls a and b of leaf extracts in different solvents. In Advances in Photosynthesis Research: Proceedings of the VIth International Congress on Photosynthesis, Brussels, Belgium, August 1–6, 1983 Volume 2 (pp. 9–12). Springer Netherlands.

21. Liopa-Tsakalidi, A., Chalikiopoulos, D., & Papasavvas, A. (2010). Effect of chitin on growth and chlorophyll content of two medicinal plants. J. Med. Plants Res, 4(7), 499–508.

22. Maleita, C. M., Curtis, R. H., Powers, S. J., & de O. Abrantes, I. M. (2012). Inoculum levels of Meloidogyne hispanica and M. javanica affect nematode reproduction, and growth of tomato genotypes. Phytopathologia Mediterranea, 566–576.

23. Martins, S. J., Pasche, J., Silva, H. A. O., Selten, G., Savastano, N., Abreu, L. M., … & Cernava, T. (2023). The use of synthetic microbial communities to improve plant health. Phytopathology®, 113(8), 1369–1379.

24. Mohindra, A., Chauhan, A., & Prabha, V. (2020). Purification and characterization of collagenase from Bacillus altitudinis. JMRR, 45.

25. Neu, A. T., Allen, E. E., & Roy, K. (2021). Defining and quantifying the core microbiome: challenges and prospects. Proceedings of the National Academy of Sciences, 118(51), e2104429118.

26. Page, A. P., Stepek, G., Winter, A. D., & Pertab, D. (2014). Enzymology of the nematode cuticle: A potential drug target?. International Journal for Parasitology: Drugs and Drug Resistance, 4(2), 133–141.

27. Pasche, J. M., Brito, J. A., Vallad, G. E., Brawner, J., Snyder, S. L., Fleming, E. A., … & Martins, S. J. (2023). Assessing the impact of successive soil cultivation on Meloidogyne enterolobii infection and soil bacterial assemblages. Plant Pathology, 72(7), 1326–1334.

28. Poudel, R., Jumpponen, A., Kennelly, M. M., Rivard, C., Gomez-Montano, L., & Garrett, K. A. (2023). Integration of Phenotypes in Microbiome Networks for Designing Synthetic Communities: a Study of Mycobiomes in the Grafted Tomato System. Applied and Environmental Microbiology, 89(6), e01843–22.

29. Poudel, R., Jumpponen, A., Schlatter, D. C., Paulitz, T. C., Gardener, B. M., Kinkel, L. L., & Garrett, K. A. (2016). Microbiome networks: a systems framework for identifying candidate microbial assemblages for disease management. Phytopathology, 106(10), 1083–1096.

30. Pichyangkura, R., & Chadchawan, S. (2015). Biostimulant activity of chitosan in horticulture. Scientia Horticulturae, 196, 49–65.

31. R Core Team. (2024). R: A language and environment for statistical computing. RFoundation for Statistical Computing. Vienna, Austria. https://www.R-project.org/

32. Raina, A., Danish, M., Khan, S., & Sheikh, H. (2020). Role of biological agents for the management of plant parasitic nematodes. Plant Pathogens (pp. 181–200). Apple Academic Press.

33. Rashidifard, M., Fourie, H., Ashrafi, S., Engelbrecht, G., Elhady, A., Daneel, M., & Claassens, S. (2022). Suppressive Effect of Soil Microbiomes Associated with Tropical Fruit Trees on Meloidogyne enterolobii. Microorganisms, 10(5), 894.

34. Riseh, R. S., Vazvani, M. G., Vatankhah, M., & Kennedy, J. F. (2024). Chitin-induced disease resistance in plants: A review. International Journal of Biological Macromolecules, 131105.

35. Sanz-Alférez, S., Mateos, B., Alvarado, R., & Sánchez, M. (2008). SAR induction in tomato plants is not effective against root-knot nematode infection. European journal of plant pathology, 120, 417–425.

36. Sauheitl, L., Glaser, B., & Weigelt, A. (2009). Uptake of intact amino acids by plants depends on soil amino acid concentrations. Environmental and Experimental Botany, 66(2), 145–152.

37. Sawaguchi, A., Ono, S., Oomura, M., Inami, K., Kumeta, Y., Honda, K., … & Saito, A. (2015). Chitosan degradation and associated changes in bacterial community structures in two contrasting soils. Soil Science and Plant Nutrition, 61(3), 471–480.

38. Schrempf, H. (2001). Recognition and degradation of chitin by streptomycetes. Antonie Van Leeuwenhoek, 79, 285–289.

39. Shahrajabian, M. H., Chaski, C., Polyzos, N., Tzortzakis, N., & Petropoulos, S. A. (2021). Sustainable agriculture systems in vegetable production using chitin and chitosan as plant biostimulants. Biomolecules, 11(6), 819.

40. Simmons, T., Caddell, D.F., Deng, S., Coleman-Derr, D. Exploring the Root Microbiome: Extracting Bacterial Community Data from the Soil, Rhizosphere, and Root Endosphere. J. Vis. Exp. (135), e57561, doi:10.3791/57561 (2018).

41. Tian, H., Riggs, R. D., & Crippen, D. L. (2000). Control of soybean cyst nematode by chitinolytic bacteria with chitin substrate. Journal of Nematology, 32(4), 370.

42. Toju, H., Peay, K. G., Yamamichi, M., Narisawa, K., Hiruma, K., Naito, K., … & Kiers, E. T. (2018). Core microbiomes for sustainable agroecosystems. Nature plants, 4(5), 247–257.

43. Tran, L. H., & Nagano, H. (2002). Isolation and characteristics of Bacillus subtilis CN2 and its collagenase production. Journal of food science, 67(3), 1184–1187.

44. Tuncsoy, B. (2021). Nematicidal activity of silver nanomaterials against plant-parasitic nematodes. Silver Nanomaterials for Agri-Food Applications (pp. 527–548). Elsevier.

45. Vieira de Carvalho Júnior, O., de Sá, A. V., Peixoto, A. R., da Paz, C. D., da Cunha e Castro, J. M., & Gava, C. A. T. (2022). Local Bacillus species as potential biocontrol agents for *Meloidogyne enterolobii* in melon (Cucumis melo L.). Biocontrol Science and Technology, 32(3), 314–328.

46. Vinolas, L. C., Healey, J. R., & Jones, D. L. (2001). Kinetics of soil microbial uptake of free amino acids. Biology and fertility of soils, 33, 67–74.

47. Volant, S., Lechat, P., Woringer, P., Motreff, L., Campagne, P., Malabat, C., … & Ghozlane, A. (2020). SHAMAN: a user-friendly website for metataxonomic analysis from raw reads to statistical analysis. BMC bioinformatics, 21, 1–15.

48. Wang, Z., Hu, X., Solanki, M. K., & Pang, F. (2023). A synthetic microbial community of plant core microbiome can be a potential biocontrol tool. Journal of Agricultural and Food Chemistry, 71(13), 5030–5041.

49. Withers, E., Hill, P. W., Chadwick, D. R., & Jones, D. L. (2020). Use of untargeted metabolomics for assessing soil quality and microbial function. Soil Biology and Biochemistry, 143, 107758.

50. Zhou, W., Wheeler, T. A., Starr, J. L., Valencia, C. U., & Sword, G. A. (2018). A fungal endophyte defensive symbiosis affects plant-nematode interactions in cotton. Plant and soil, 422, 251–266.

51. Zhou, X., Wang, J., Liu, F., Liang, J., Zhao, P., Tsui, C. K., & Cai, L. (2022). Cross-kingdom synthetic microbiota supports tomato suppression of Fusarium wilt disease. Nature communications, 13(1), 7890.

52. Zhou, Z., Gao, T., Van Zwieten, L., Zhu, Q., Yan, T., Xue, J., & Wu, Y. (2019). Soil microbial community structure shifts induced by biochar and biochar-based fertilizer amendment to Karst calcareous soil. Soil Science Society of America Journal, 83(2), 398–408

53. Zitouni, M., Viens, P., Ghinet, M. G., & Brzezinski, R. (2017). Diversity of family GH46 chitosanases in Kitasatospora setae KM-6054. Applied microbiology and biotechnology,101, 7877–7888.

